# Assessment of Meridic Gel Larval Diet and Meridic Adult Diet for Mass Rearing of *Bactrocera dorsalis* (Diptera: Tephritidae)

**DOI:** 10.1101/2025.03.10.642425

**Authors:** Mahfuza Momen, Md. Shahjalal, Md. Ashikur Rahman, Md. Aftab Hossain, Md. Kamruzzaman Munshi, Kajla Seheli

## Abstract

The oriental fruit fly, *Bactrocera dorsalis* (Diptera: Tephritidae), is a widespread pest in Bangladesh. Sterile Insect Technique (SIT) offers a solution for effectively suppressing this fruit fly species. However, SIT involves mass rearing of fruit fly species in a laboratory where a standardized artificial rearing diet is crucial for ensuring uniform growth, development, and reproduction. In this study, we assessed efficacy of a new formulated gel-based meridic larval diet as well as protein and carbohydrate rich adult diets for the rearing of *B. dorsalis* in a laboratory condition. Proximate analysis was conducted for our formulated rearing diets to determine the content of moisture, protein, fat, carbohydrate, and ash. Several key biological parameters, including egg hatching rate, pupation rate, pupal weight, adult emergence, adult growth, sex ratio, and flight capacity, were assessed. Statistical analysis using Tukey box plots revealed a significant improvement in adult body parameters and an increase in longevity for our formulated diets. In addition, our study presents survival analysis using non-parametric Kaplan–Meier estimator and Weibull parametric model. This study shows that integrating our formulated diets into the mass rearing process of *B. dorsalis* in a laboratory could significantly improve the production of larger and healthier flies on a large scale.

## 1. Introduction

*Bactrocera dorsalis*, commonly known as oriental fruit fly, is a highly destructive dominant pest of fruits and vegetables in Bangladesh (Leblanc *et al*., 2019; Momen et al., 2024; Vargas et al., 2015). *B. dorsalis* females lay their eggs in the pulp of a wide range of hosts, leading to a significant crop damage as the larvae hatch and begin feeding on fruits. After going through three stages of development, the larvae eventually leave the fruits to pupate in the soil from where adults emerge. Again, they will continue the cycle by laying eggs into new fruits, ultimately leading to further losses to farmers (Jaffar *et al*., 2023). The implementation of proper control measures for this pest is crucial in order to mitigate economic loss and to ensure the protection of agricultural yields. Among various approaches of effective pest management strategies, the Sterile Insect Technique (SIT) offers an effective and sustainable approach for controlling of *B. dorsalis* (Steiner *et al*., 1970; Vargas *et al*., 2015).

### Larval diet

Mass rearing of *B. dorsalis* larvae in a laboratory is one of the crucial primary steps for a successful implementation of SIT. In a laboratory, a larval diet creates an environment for larvae to grow and feed before they pupate. By ensuring optimal nutrition and conditions for larval development, we can increase the production of sexually competitive adults for a successful SIT application. Therefore, the implementation of appropriate cost-effective larval diets that ensure optimal larval development and lead to mass production of healthy competitive males, is an integral part for successful SIT programs (Wang *et al*., 2023). However, suitable larval and adult diets for mass rearing of this fruit fly in a laboratory can be an expensive and challenging process (Parker, 2005). The primary reason for this is the absence of a standard rearing diet that can be made from locally available and affordable ingredients.

Primarily, whole host fruits like papaya, banana, guava etc. were used to establish the colony in many laboratories on a small scale (Vargas *et al*., 1996). However, fruits are typically seasonal, which limits the availability of fresh fruits in consistent quantities needed for a larval diet. Furthermore, use of natural hosts for laboratory rearing of fruit flies introduces inconsistencies and variability in quality (Newman *et al*., 2021). Therefore, alternative larval diets are required to overcome these limitations.

A major advancement in the laboratory rearing tephritid fruit fly larvae was the incorporation of dehydrated plant materials (e.g., carrot powder) and dried yeast into their diets. This addition of food significantly improved larval rearing and resulted in higher pupal yields (Dominguez Gordillo, 1999; Mitchell *et al*., 1965). Over the years, several artificial larval diets have been developed, used, and evaluated for mass rearing of *B. dorsalis* (Chang, 2009b; Chang et al., 2006; Hou et al., 2020; Khan et *al*., 2019). Mass production of *B. dorsalis* is usually based on a diet that includes water, sugar and yeast as nutrients, and grain powder as a bulking agent. However, different types of bulking agents (e.g., corn cob powder, bagasse, wheat bran, etc.) have major drawbacks, such as excessive heat generation due to microbial activity and a tendency for mycotoxin contamination, negatively impacting growth and survival. Another significant drawback of this formulation is the substantial waste generation due to the use of bulking agents. This undesirable waste, which remains useless as well as negatively impacts the rearing environment, also increases production cost. Therefore, researchers are in search of a diet with a new formulation that would ensure sustainable mass rearing by reducing bulking agents.

Gelling agents, which can prevent undesirable reactions among ingredients of a diet, are potential ingredients that can be used in larval diets to convert high-water content mixtures into semi-solid gels with a homogeneous distribution of ingredients. For instance, agar has exceptional water retention properties, capable of holding a high percentage of water in its gel form (Xiao *et al*., 2019). This attribute of agar helps to reduce desiccation as well as maintain the larval diet texture (Pascacio-Villafán *et al*., 2020). As a result, gel-based diets emerged as a promising alternative for larval rearing. In our laboratory, a gel-based larval diet was adopted for rearing *B. dorsalis* (Chang, 2009b; Khan *et al*., 2019). However, for scaling up mass-production of *B. dorsalis* from small-scale laboratory rearing, there is a search for a more efficacious and cost-effective larval diet formulation. A very limited number of research works were reported on the larval gel diets (Hou *et al*., 2020; Khan *et al*., 2019; Pascacio-Villafán *et al*., 2020; Pásková, 2007; Prekas *et al*., 2023). Previous research by Khan *et al*. explored a gel-based larval diet to optimize the rearing of a local wild *B. dorsalis* population (Chang, 2009b; Khan *et al*., 2019). However, in the previous study, a limited set of biological quality parameters was examined without analyzing the nutrient contents of the diet. In addition to that various parameters have interdependent relationships with one another. Therefore, there is a need for more comprehensive studies where relevant parameters are studied together to get a more detailed understanding of the efficacy of diet formulation.

### Adult diet

Healthy and vigorous fruit flies are essential for research and SIT programs. A proper diet for adult rearing is crucial for maintaining the quality of adult fruit flies. However, mass rearing of the adult stage presents subsequent challenges (Canale *et al*., 2015; FAO/IAEA/USDA, 2019; Shelly, 2017; Sunday & Samira, 2011). While newly emerged adults can survive on sugar and water, dietary protein is also essential for gonad maturation and male mating performance. Therefore, understanding the nutritional requirements and dietary preferences of adult Tephritidae is extremely important for developing effective management strategies as well as improving the quality of mass-reared individuals for pest control programs. Careful consideration of the sources and balance of key nutrients like proteins, carbohydrates, vitamins, and lipids is essential for formulating an efficient artificial diet for adult *B. dorsalis*. Previous studies have shown that diets containing yeast or yeast derivatives enhance egg production and overall fitness of various Tephritidae species (Chang, 2009a; Goane *et al*., 2018). A common formulation includes a mixture of sugar and yeast. This combination provides the essential nutrients for adult flies, promoting mating behavior and longevity (Shelly & Edu, 2007). Containing vitamins in the diet is also important for reproductive health and stress resistance as well as overall fitness of adult fruit flies. The composition of the adult diet directly affects sexual maturation rates, body weight, and mating success. Diets that are rich in vitamins and proteins can improve longevity under stressful conditions, which is critical for maintaining a healthy population during mass rearing.

An ideal diet should maintain a consistent composition that prevents fermentation as well as eliminates the necessity for additional supporting substrates and minimizes waste during the rearing process. Such a diet simplifies rearing procedures and ensures a consistent nutritional intake, leading to more standardized insect quality. Several formulations of artificial adult diet have been developed using low-cost ingredients for different tephritid species (Barry *et al*., 2007; Canale *et al*., 2015; Goane *et al*., 2018; Tsitsipis & Kontos, 1983). However, advancement in the diet formulations is still necessary to optimize it further, leading to improved stability and reduced fermentation, especially in the subtropical monsoon climate of Bangladesh.

Despite progress in adult diet development, limited information is available regarding the effect of both dry and liquid formulations of adult diets on the lifespan as well as overall fitness of adult oriental fruit flies. Further research that takes into account various parameters simultaneously is essential for understanding the long-term impacts of adult diets on the lifespan and quality of adult *B. dorsalis*. To assess the quality and longevity of *B. dorsalis* adults under laboratory rearing conditions, in this study we conducted a series of experiments using two types of adult diet formulations (dry and liquid diet).

In this work, we present a comprehensive assessment and comparison of the effectiveness of our new formulated gel-based meridic diet for larvae along with yeast-based diets for adult fruit flies. Our study presenting Tukey box plots demonstrated a significant improvement in adult body parameters and longevity while using the new formulated diets. In addition, we preformed survival analysis using non-parametric Kaplan–Meier estimator and Weibull parametric model. The findings in this study will contribute to the optimization of rearing techniques ensuring the sustainability and efficiency of the mass production of this fruit fly species.

## 2. Material and Methods

This study was conducted at Insect Biotechnology Division laboratory of Institute of Food and Radiation Biology, Atomic Energy Research Establishment, Bangladesh. The fruit fly species, *B. dorsalis*, used in this experiment was obtained from a laboratory colony established from a local pest population. The insectary was maintained at 27±1 °C, 70±5% relative humidity (RH), with a 14-h light:10-h dark photoperiod. All the procedures used in this study are summarized below.

### 2.1. Larval diet formulation

In this study, the new formulated larval diet used for rearing *B. dorsalis* larvae was gel-based. To prepare this larval diet, all the ingredients (sugar, preheated baking yeast, soy protein, soy bran, sodium benzoate, citric acid, agar-agar and distilled water) were mixed well (see Table 1).

**Table 1.**
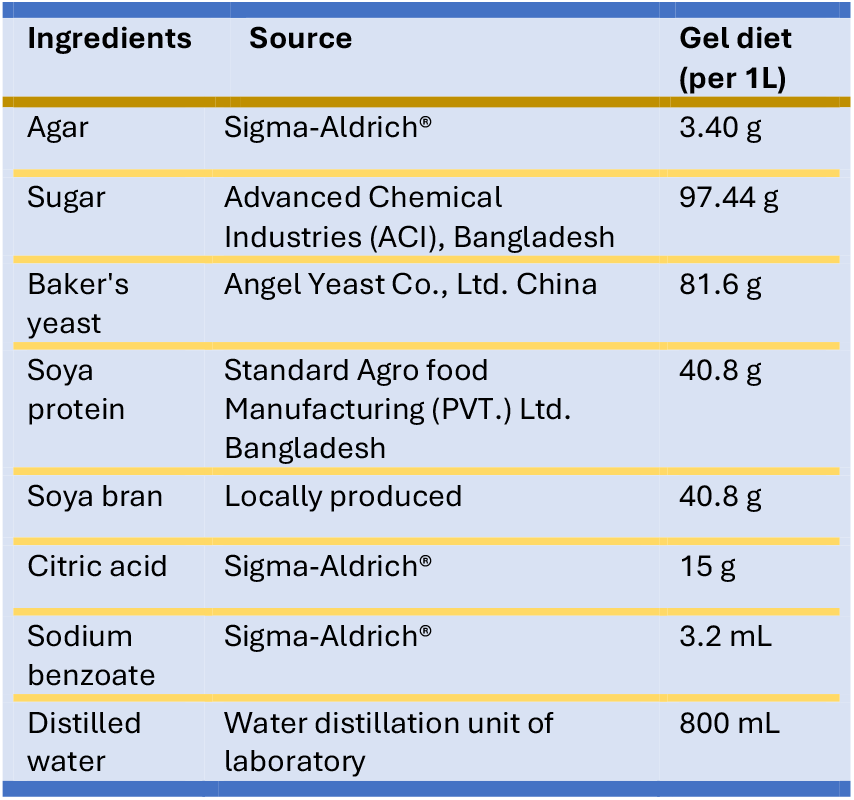
Formulation of gel-based larval diet for rearing *Bactrocera dorsalis*.

| Ingredients | Source | Gel diet (per 1L) |
| --- | --- | --- |
| Agar | Sigma-Aldrich® | 3.40 g |
| Sugar | Advanced Chemical Industries (ACI), Bangladesh | 97.44 g |
| Baker's yeast | Angel Yeast Co., Ltd. China | 81.6 g |
| Soya protein | Standard Agro food Manufacturing (PVT.) Ltd. Bangladesh | 40.8 g |
| Soya bran | Locally produced | 40.8 g |
| Citric acid | Sigma-Aldrich® | 15 g |
| Sodium benzoate | Sigma-Aldrich® | 3.2 mL |
| Distilled water | Water distillation unit of laboratory | 800 mL |

The mixture was carefully heated until it reached a vigorous boil, and then the boiling temperature was maintained for 5 minutes to ensure thorough heating. Aliquots of 333 mL of the hot gel-based diet were then dispensed into individual larval rearing trays (14 cm × 20 cm) and left to solidify at room temperature for 2-3 hours.

Following the preparation of the larval diet, we collected *B. dorsalis* eggs using oviposition devices. After allowing the flies to oviposit for 1 hour, eggs were collected and rinsed from the oviposition device into a beaker. After that 0.5 mL eggs were then placed on the pieces of moistened soft non-woven felt fabric (1 mm × 2 cm × 4 cm) and kept on top of the diet surface within 20 minutes of collection.

After adding eggs to the diet, the rearing trays were transferred into larger plastic containers with dimensions of 37 cm in length, 27 cm in width, and 10 cm in height. The containers were securely sealed with a lid that had a ventilation opening measuring 15 cm by 10 cm. The opening was covered by a mesh to ensure adequate airflow within the containers. All the rearing trays were kept within these plastic boxes at a constant temperature of 27±1 °C, until the larvae began to exit the diet to pupate. Before the larvae could jump, the rearing trays were transferred to a similar box filled with sawdust at the bottom (Figure 1 (a)). In this pupation box, sawdust was used as a pupation substrate. The pupae were collected by sieving the sawdust and then placed in petri dishes. Finally, the pupae were placed in adult rearing cages (31 cm × 23 cm × 22 cm) for adult emergence. Flies were then kept at 27±1 °C, 70±5% relative humidity (RH), with a 14-h light:10-h dark photoperiod in adult rearing cages.

**Figure 1.**
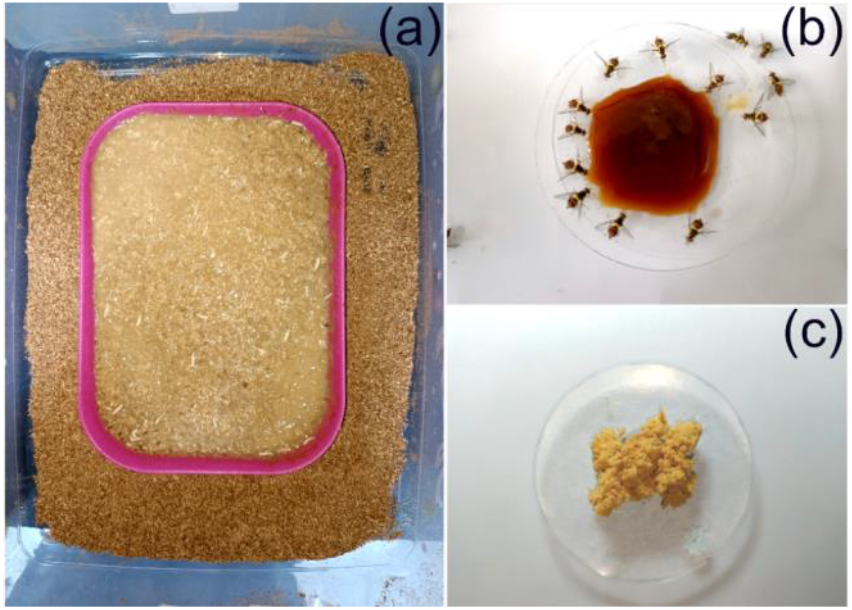
Artificial diets for mass rearing of *Bactrocera dorsalis*: (a) gel-based meridic larval diet, (b) liquid adult diet, (c) dry adult diet.

### 2.2. Adult diet formulation

All ingredients used in the formulation of adult diets for rearing *B. dorsalis* are shown in Table 2. The adult flies in this study were provided with unlimited access to two different yeast-based diets: a protein-rich dry diet and a liquid diet (Table 2). The yeast used in the diets not only contained protein but also provided essential B-complex vitamins, sodium chloride, lipids, and minerals. In addition, the casein used in dry adult diet was another source of protein. The source of carbohydrates was sugar. This formulation ensured that the flies received a well-rounded nutritional intake. Water was supplied using a cotton wick positioned in a conical flask filled with water.

**Table 2.** Formulations of adult diets for rearing *Bactrocera dorsalis*.

| Ingredients | Source | Dry diet (per 1kg) | Liquid diet (per 1kg) |
| --- | --- | --- | --- |
| Yeast extract | Sigma-Aldrich® | 250 g | - |
| Casein | Sigma-Aldrich® | 250 g | - |
| Sugar | Advanced Chemical Industries (ACI) Foods Ltd. Bangladesh | 500 g | 300 g |
| Baker's yeast (Incubated in oven at 47 °C) | Angel Yeast Co., Ltd. China | - | 100 g |
| Distilled water | Water distillation unit of laboratory | - | 600 mL |
| **Note:** Incubate the entire mixture of adult liquid diet in an oven at 47 °C for up to 5 days. |  |  |  |

The dry adult diet (Figure 1 (c)) used in the experiment was prepared by directly mixing all the ingredients: casein, yeast extract, and sugar in the ratio of 1:1:2. In this dry adult diet formulation, casein has a water-binding capacity from the air due to its structure. This water binding capacity of casein allows it to hold a significant amount of water within its protein network. In addition to the dry adult diet, another long-lasting liquid diet was also provided (Figure 1 (b)).

This adult liquid diet consisted of distilled water, sugar, and baker’s yeast (*Saccharomyces cerevisiae*) (Table 2). The initial liquid diet mixture was incubated in an oven at 47 °C for five days until a thick syrup with high viscosity was formed. Both dry and liquid diets were offered to adults in separate watch glasses, allowing for ad-libitum consumption throughout the duration of the experiment.

### 2.3. Nutrient analysis of adult and larval diets

Before using our new formulated adult and larval diets as rearing diets, a nutrient analysis was conducted using proximate analysis method at Food Safety and Quality Analysis Division of the Institute of Food and Radiation Biology, Atomic Energy Research Establishment, Bangladesh.

### 2.4. Flight test

The assay was carried out according to FAO/IAEA/USDA quality control manual (FAO/IAEA/USDA, 2019) with minor modifications. To evaluate the fitness of adult flies, we used a device made from a black PVC pipe. The pipe had a diameter of 9 cm and a height of 10 cm. The bottom of the pipe was sealed with a black paper-lined petri dish (9 cm in diameter). Prior to the flight test, the inner walls of the PVC pipe were dusted with odorless talcum powder to stop the flies from walking out. The percentage of flies that were able to fly was estimated using the following equation (Chang, 2004):

Percentage of fliers

= Number of total pupae −

((Number of not emerged pupa

+Number of partially emerged pupae

+Number of deformed pup

+Number of nonfliers)/(Number of total pupae)) × 100

### 2.5. Morphometry

Different physical parameters of adult *B. dorsalis* including body length, wing length, thorax width, etc. were measured. In addition, pupal size and weight were also measured for *B. dorsalis* reared on larval and adult diet under laboratory conditions (see Table 5, Table 6 and Figure 2). To compare biological parameters, wild *B. dorsalis* adults were collected. The wild males were collected from the local population using methyl eugenol baited traps. On the other hand, the wild females were collected from six types of infested fruits (guava, wax apple, slow match tree, mango, and tropical almond).

**Figure 2.**
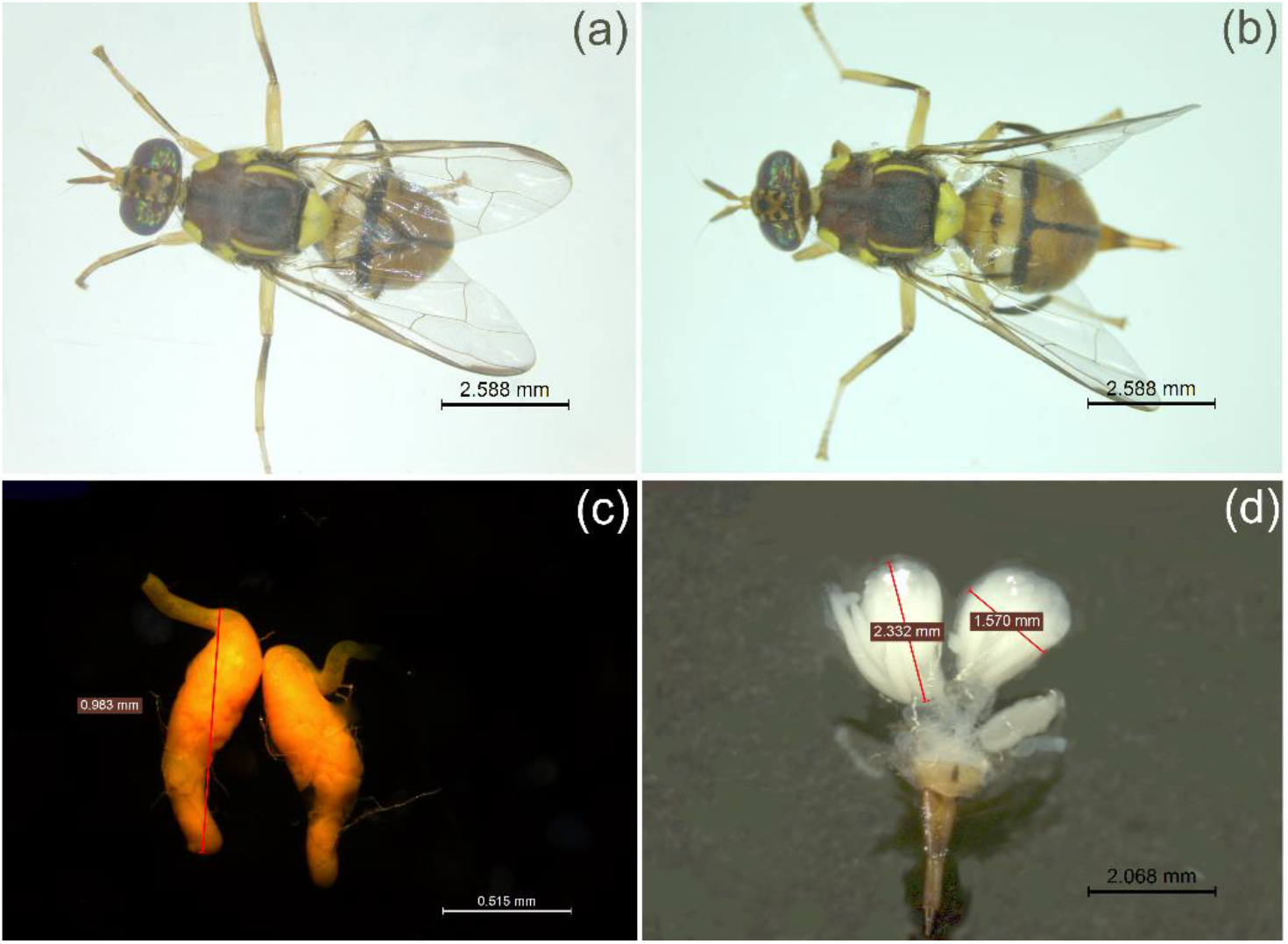
Morphological details of *Bactrocera dorsalis*: (a) adult male, (c) testes, (b) adult female, and (d) ovaries.

Furthermore, morphometry of ovary and testes were also observed (Figure 2) under a stereomicroscope (Leica S8 APO Stereo Microscope with Leica DMC2900 digital camera) where the necessary parameters like ovary length and testes length were measured. First, sexually mature male and female flies (15 days old) were collected from adult rearing cages and the collected flies were killed in 70% ethanol before dissection. Next, the reproductive systems were extracted from the lower abdomen of adult flies (Figure 2). The reproductive systems were then carefully removed from the surrounding tissues following the dissection procedure described by Chou *et al*. (Chou *et al*., 2012). Similar measurements were also taken for the collected wild male and female specimens, excluding the ovary of wild females.

### 2.6. Survival analysis

Adult flies were placed in rearing cages immediately after their emergence to evaluate their longevity and survival. In one experiment, the flies were given unlimited access to the two diets (dry and liquid adult diets as mentioned in the 2.2) where the two diets were provided in separate watch glasses (Figure 1 (b, c)). In another experiment, longevity and survival under stress were assessed by providing only distilled water.

For adult survival analysis, both male and female data were calculated separately as their longevity was different. In this study, survival analyses were performed using two different widely used survival analysis methods (non-parametric Kaplan–Meier estimator and Weibull parametric model) by biologists. The survival curves of both males and females those kept under similar conditions were compared with each other. In this study, we recorded the mortality rates on a daily basis.

## 3. Results and Discussion

### 3.1. Nutrient analysis of larval and adult diets

Analyses of nutrient composition of gel-based larval diet and adult diets (dry and liquid) used for *B. dorsalis* rearing (Figure 1) revealed actual ratio of moisture, protein, fat, carbohydrate, and ash content. Table 3 shows the mean values of these nutrient contents in percentage with standard deviation (SD). Initial pH measurements, as shown in Table 4, indicated that all the formulated diets were slightly to moderately acidic. These pH levels of diets are helpful to suppress harmful microbial growth, which is beneficial for fruit fly health.

**Table 3.** Nutrient contents of larval and adult diets used for rearing *Bactrocera dorsalis*.

| Nutrient | Gel-based larval diet<br>(Mean ± SD) % | Dry adult diet<br>(Mean ± SD) % | Liquid adult diet<br>(Mean ± SD) % |
| --- | --- | --- | --- |
| Moisture | 72.22 ± 0.39 | 8.67 ± 0.33 | 20.78 ± 0.69 |
| Protein | 5.83 ± 0.20 | 35.82 ± 0.73 | 7.99 ± 0.10 |
| Fat | 2.48 ± 0.01 | 5.52 ± 0.09 | 1.04 ± 0.01 |
| Carbohydrate | 18.73 ± 0.15 | 47.48 ± 0.45 | 69.41 ± 0.56 |
| Ash | 0.73 ± 0.03 | 2.52 ± 0.03 | 0.78 ± 0.05 |
| **Note:** The analysis of the larval diet was conducted after gelation process. |  |  |  |

**Table 4.** Initial acidity or alkalinity of larval and adult diets used for rearing *Bactrocera dorsalis*.

| Category of diet | Acidity or alkalinity (pH)<br>(Mean ± SD) |
| --- | --- |
| Gel-based larval diet | 4.66 ± 0.006 |
| Dry adult diet | 6.56 ± 0.01 |
| Liquid adult diet | 5.17 ± 0.01 |
| **Note:** The pH analysis was conducted after diet preparation was completed. |  |

**Table 5.**
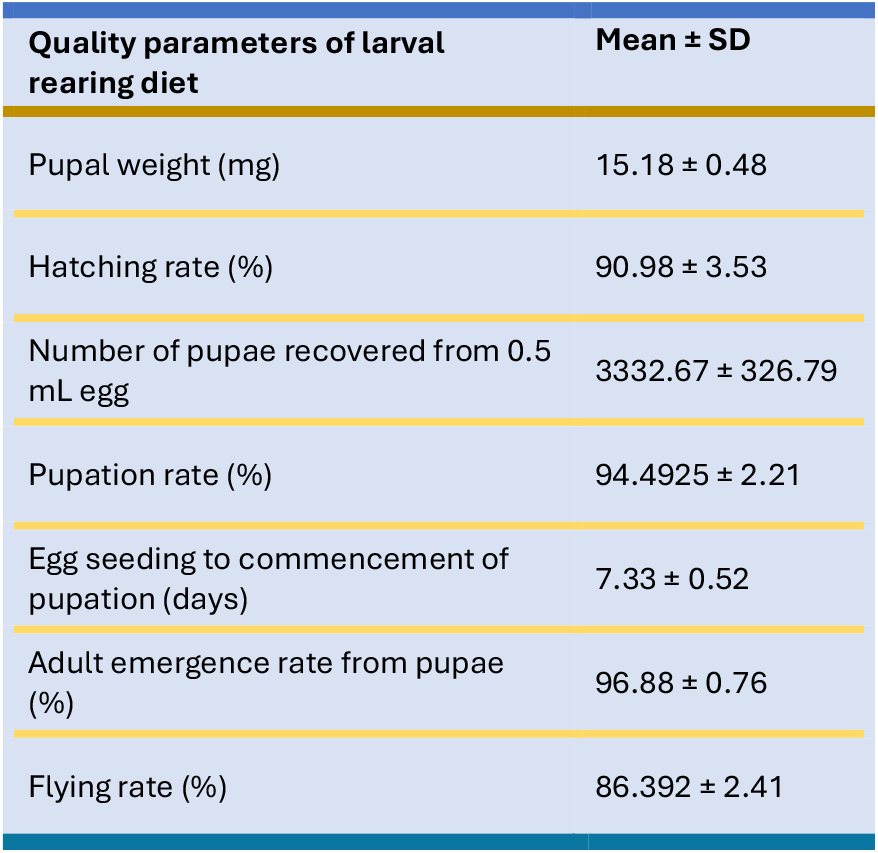
Quality parameters of larval rearing diet for *Bactrocera dorsalis*.

**Table 6.**
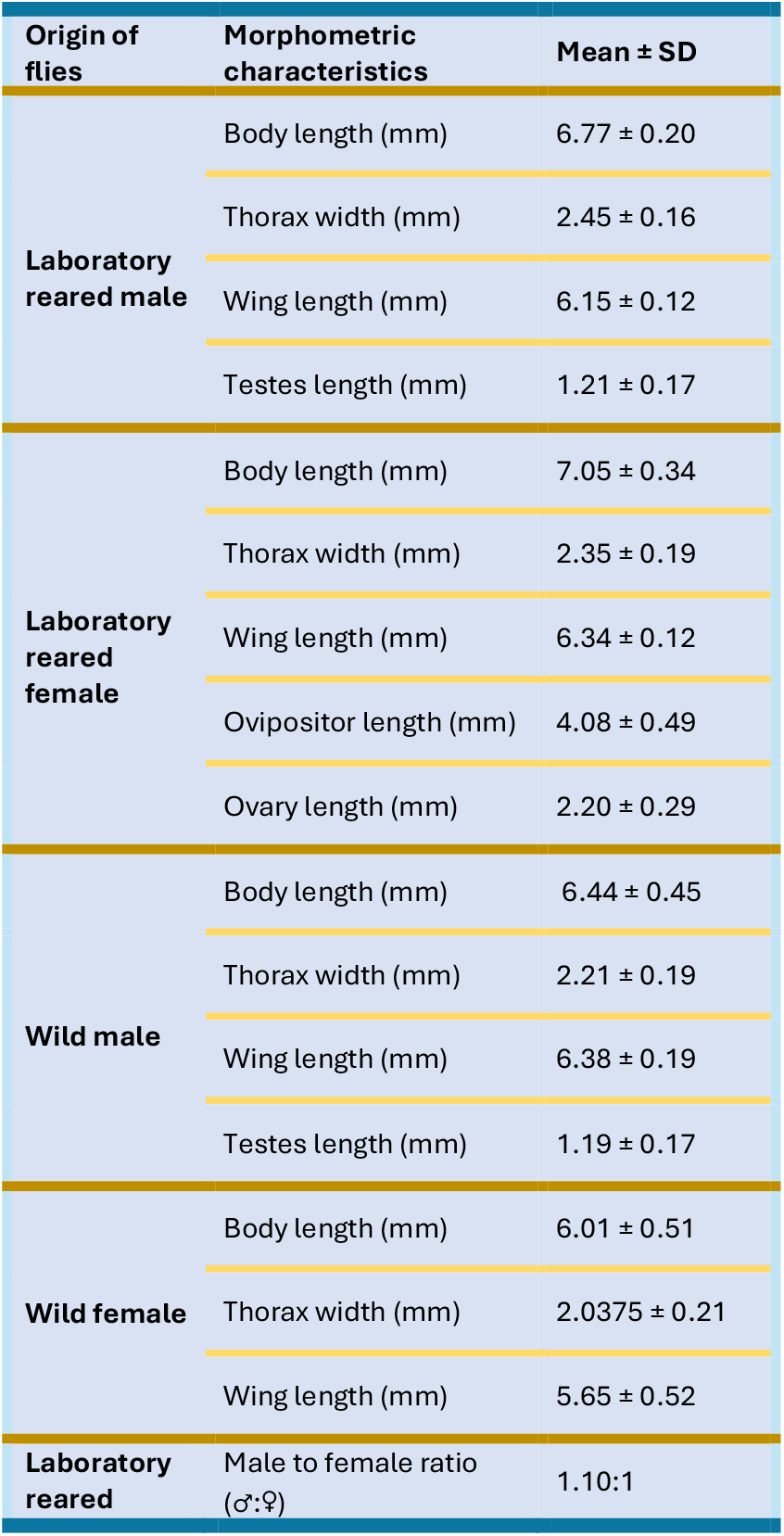
Quality parameters of adult *Bactrocera dorsalis*.

The initial acidity (pH) for our formulated larval diet was (pH 4.66±0.006), which is higher compared to the diet described by Mahfuza *et al*. (pH in the range of 3.5 to 4) (Khan *et al*., 2019). As compared to the diet formulation described by Mahfuza *et al*., we used different composition ratio of ingredients in our larval diet formulation. As no bacteria was used as probiotics in our diet formulation, this formulation involves lower production cost.

### 3.2. Influence on egg hatching rate

The egg hatching rate observed in this larval diet was 90.98±3.53%, which is very close with the value 90.92±0.9% reported by C. Chang for the larval liquid diet (Chang, 2009a). The hatching rate of *B. dorsalis* in an artificial diet (wheat bran media) reported by Suhana *et al*. was 84.3%, which is lower compared to our larval diet (Yusof *et al*., 2019). According to previous reports by C. Chang and Pieterse *et al*., the observed hatching rate in this study indicates that our formulated diet is suitable for mass rearing (Chang, 2009a; Pieterse *et al*., 2019).

### 3.3. Influence on pupal weight

Different quality parameters (including pupal size, weight, and adult wing size) of *B. dorsalis* were also observed (Table 5 and Table 6) for the agar-based larval diet.

The average weight of the pupae was 15.18±0.48 mg, as shown in Table 5. According to the IAEA’s mass rearing quality control guidelines for tephritid flies (FAO/IAEA/USDA, 2019), for SIT programs the required minimum pupal weights is 12.30 mg, and acceptable mean pupal weights is 12.90 mg. The pupae reared on our diet exceeded the required average pupal weight by about 17.67% and minimum pupal weight by about 19.51%.

### 3.4. Influence on adult emergence rate

In our experiments, we observed an adult emergence rate of 96.88±1.56% (Table 5), which exceeds the International Atomic Energy Agency (IAEA) standards for mass rearing (FAO/IAEA/USDA, 2019). This adult emergence rate (pupa-to-adult fly development) exceeded both the minimum threshold of 82% as well as the acceptable threshold of 90%. The adult emergence rate of *B. dorsalis* reared on our diet exceeded the required rate by about 7.64%. The observed rate is also comparable to the 97.33% eclosion rate for *B. dorsalis* reported by Chen *et al*. (Chen *et al*., 2024). We collected a sample of 923 adult individuals to assess the adult sex ratio. The analysis revealed a male-to-female ratio of 1.10:1, indicating a slight male dominance (see Table 6). These results demonstrate that our formulated diet meets the necessary quality control standards for mass rearing of *B. dorsalis* suggested by IAEA.

### 3.5. Influence on body and wing size

Figure 2 shows morphological details of *B. dorsalis*. Different morphometric characteristics of the adults captured from wild environment as well as our laboratory reared adults fed with our formulated diet were measured (see Table 6). The Tukey box plots in Figure 3 show comparison of body length of laboratory reared and local wild *B. dorsalis* population. It was observed that laboratory-reared *B. dorsalis* adults, both male and female, were significantly larger than their wild counterparts.

**Figure 3.**
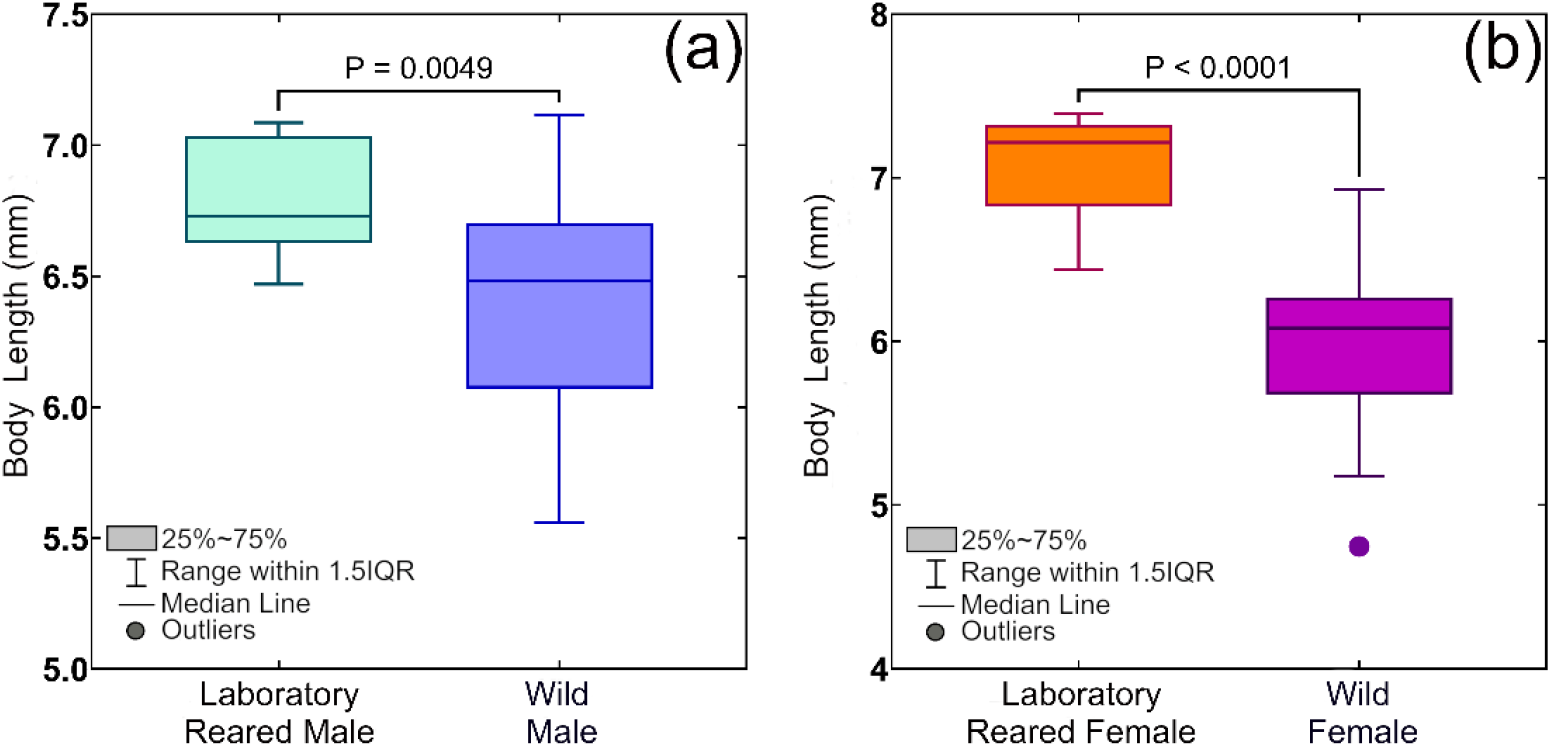
Comparison of body length of *Bactrocera dorsalis* for laboratory reared and local wild population using Tukey box plots: (a) adult males, and (b) adult females.

In addition, while comparing wing size, we observed that adult male flies had a wing size of 6.15±0.12 mm, whereas females had a slightly larger wing size of 6.34±0.12 mm (see Table 6). Previously, 6.00 mm wing size of *B. dorsalis* was reported by Drew *et al*. (Drew *et al*., 2008). The nutritional content of our formulated diet resulted in larger wings and body size, which is crucial for successful mating and population growth in laboratory conditions.

Previous research indicated that females, irrespective of their size, exhibited a preference for mating with larger males, suggesting potential advantages associated with choosing larger mates (Anjos-Duarte *et al*., 2011). Furthermore, larger males have been proven to exhibit greater pre- and post-copulatory reproductive success (Bangham *et al*., 2002). These findings suggest that the current larval-rearing diet is adequate for promoting optimal growth and overall fitness of adult flies.

### 3.6. Influence on flight ability

The ability of adult *B. dorsalis* to fly is crucial for the survival and reproductive success of their population. In our study, flight ability of adults was 86.392±2.41%, as shown in Table 5. The IAEA mass rearing manual states that *B. dorsalis* adult flies that can be employed in SIT programs should have at least 75% flight ability, whereas the ideal level of flight ability is 83% (FAO/IAEA/USDA, 2019). The measured flight ability in our study meets the standard requirements for mass rearing.

### 3.7. Influence on gonad development

Adult tephritid fruit flies require dietary protein for sexual maturation and gonad development (Jácome *et al*., 1999; Papanastasiou *et al*., 2019). Therefore, the adult flies were provided with unlimited access to two different yeast-based diets: a protein-rich dry diet and a liquid diet. In this study, we used the ovary and testes development (see Table 6) as the measurement parameters to evaluate the adult-rearing diet.

In our study, we observed that the mean ovary length of 15-day-old gravid females was 2.20±0.29 mm (see Table 6), which is larger than the previously reported ovary length of 1.8±0.07 mm (Chou *et al*., 2012).

Furthermore, we evaluated the correlation between male body size and testis size in both laboratory-reared and wild *B. dorsalis*. Figure 4 (b) shows a simple linear regression analysis which demonstrates a weak positive correlation between body size and testes size in wild males (r = 0.232, p = 0.4919).

**Figure 4.**
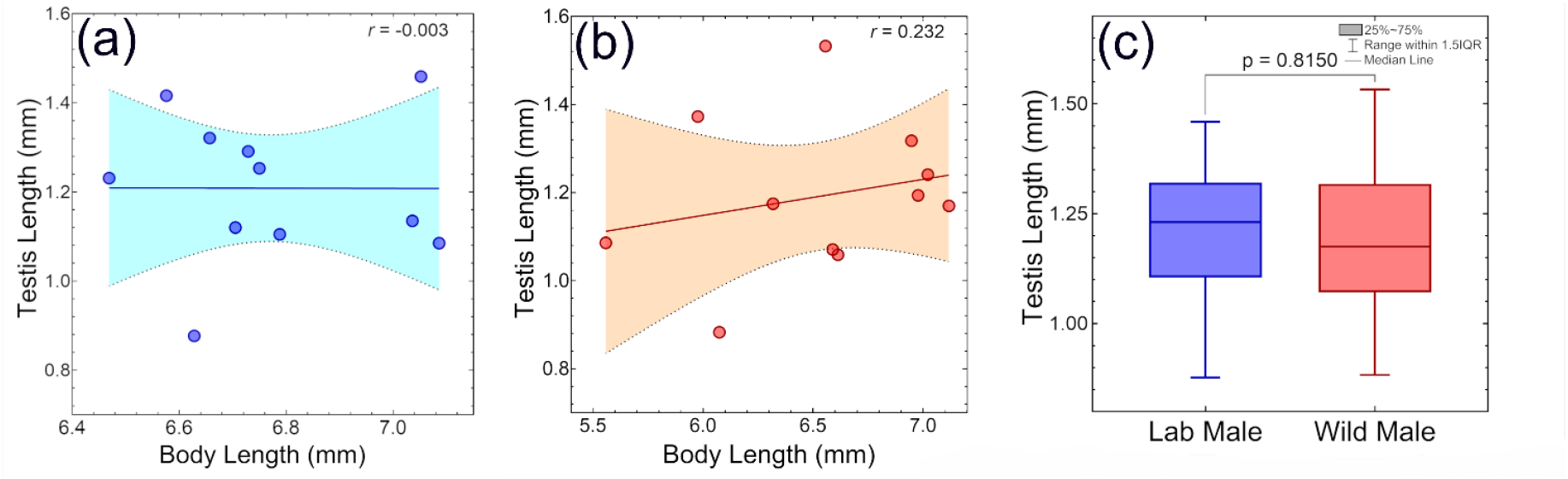
Correlation between male body size and testis size of *Bactrocera dorsalis*: (a) laboratory-reared *B. dorsalis*, (b) wild *B. dorsalis*. The color shadows represent 95% confidence interval. Figure (c) shows comparison of testis size between laboratory-reared and wild male *B. dorsalis*.

A similar size difference in testes between large and small males has also been reported for walnut flies (*Rhagoletis juglandis*) (Carsten-Conner *et al*., 2010). In the research work by Carsten-Conner *et al*. (Carsten-Conner *et al*., 2010), it was indicated that larval environment significantly influences testes size in male flies. However, this relationship was not observed in laboratory-reared males (r = -0.003, p = 0.9933) (see Figure 4(a)). This difference may be attributed to the more uniform growth observed in lab-reared males, with body lengths ranging from 6.47 mm to 7.09 mm. In contrast, wild males exhibited a wider range of body lengths (5.56 mm to 7.12 mm) (see Figure 3 (a)), possibly reflecting the influence of variable natural resources and environmental conditions on their growth and development. According to this study, the more uniform growth conditions in the laboratory may account for the lack of this association in laboratory reared individuals. The results in this study were consistent with the prediction that body size of males does not always reflect the testes size in laboratory condition on *Drosophila melanogaster* (Bangham *et al*., 2002). Furthermore, while comparing testes size of wild and lab-reared male, we found that the mean testes size for wild males was 1.19±0.18 mm (see Table 6), whereas the lab-reared males had a mean testes size of 1.208±0.1664 mm. An unpaired t-test (Figure 4 (c)) revealed that there is no statistically significant difference in testes size between the two groups (t = 0.2371, df = 20, p = 0.8150). However, previous research showed that larger male Tephritidae transfer much greater quantities of spermatozoa during mating (Msaad Guerfali & Chevrier, 2020). This competitive advantage can maximize reproductive success of laboratory reared larger males.

### 3.8. Influence on Longevity and Survival analysis

Figure 5 shows a change in longevity between male and female *B. dorsalis* under different dietary conditions. Females exhibited greater vulnerability to dietary restrictions, particularly protein and carbohydrate deprivation, as compared to their male counterparts.

**Figure 5.**
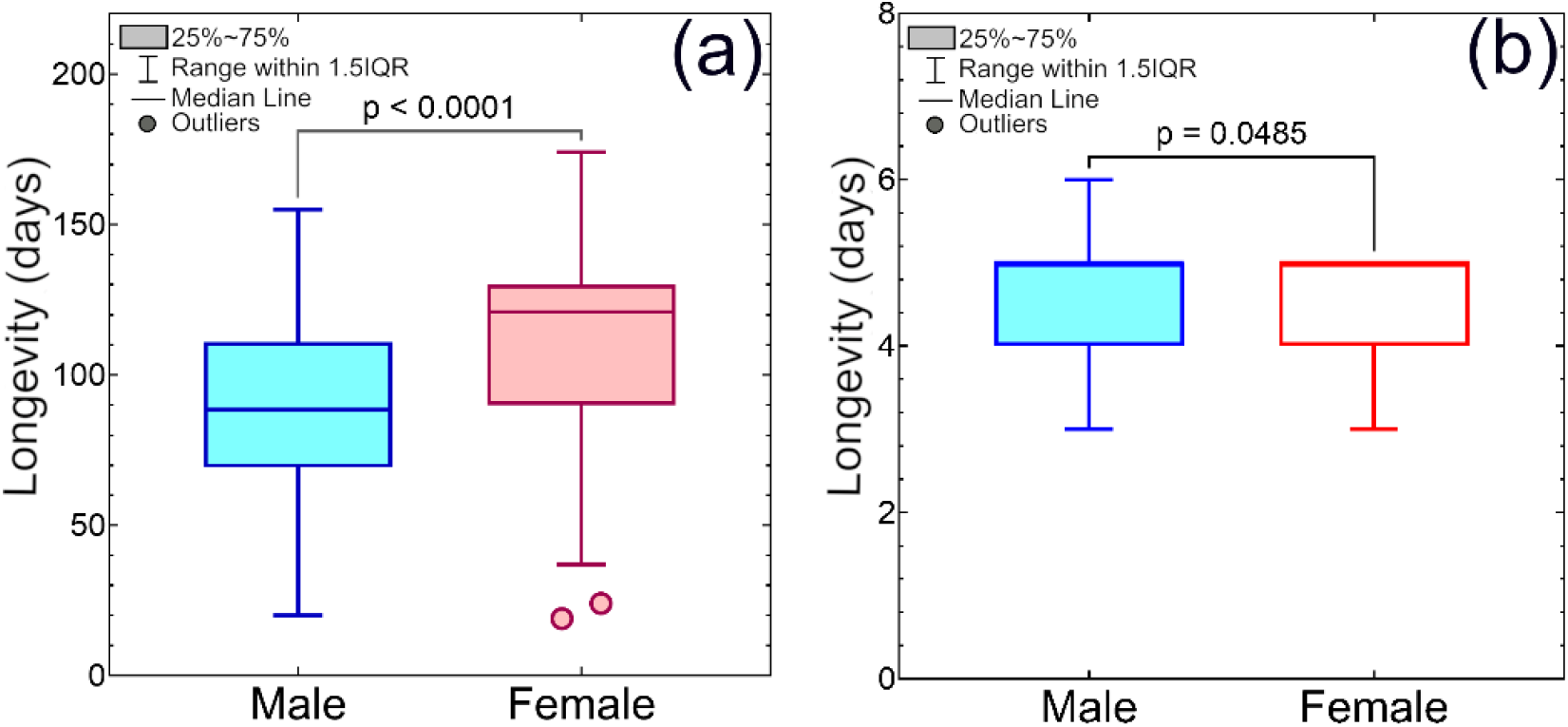
Comparison of longevity of adult male and female *Bactrocera dorsalis* using Tukey box plots: (a) Longevity of fed with carbohydrate and protein-rich food supply, (b) only distilled water supply. The median line coincides with the upper level of the box in figure (b).

Furthermore, the survival analyses (Figure 6 and Figure 7), using non-parametric Kaplan–Meier estimator and Weibull parametric model, were conducted separately for each sex for observing differences in longevity. The survival curves obtained from the survival analyses revealed an age-dependent mortality pattern, characterized by an initial increase and followed by a plateau (Figure 6 and Figure 7).

**Figure 6.**
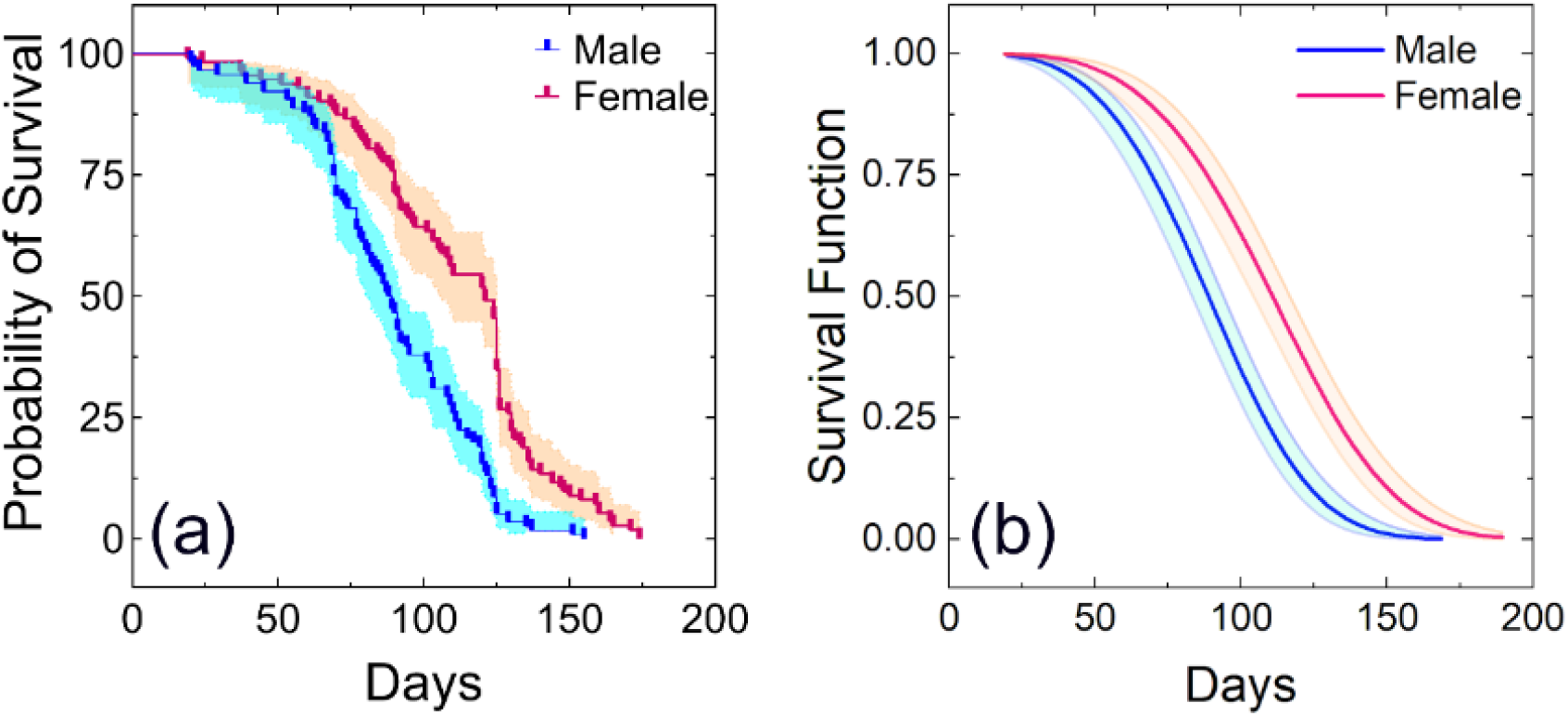
Survival curves of males (n = 116) and females (n = 112) of adult *Bactrocera dorsalis* fed with a carbohydrate and protein-rich food supply: (a) Survival curves using Kaplan–Meier estimator, (b) Survival curves using Weibull parametric model. The color shadows over the survival curves, shown in figure (a) and (b), represent the 95% confidence interval.

**Figure 7.**
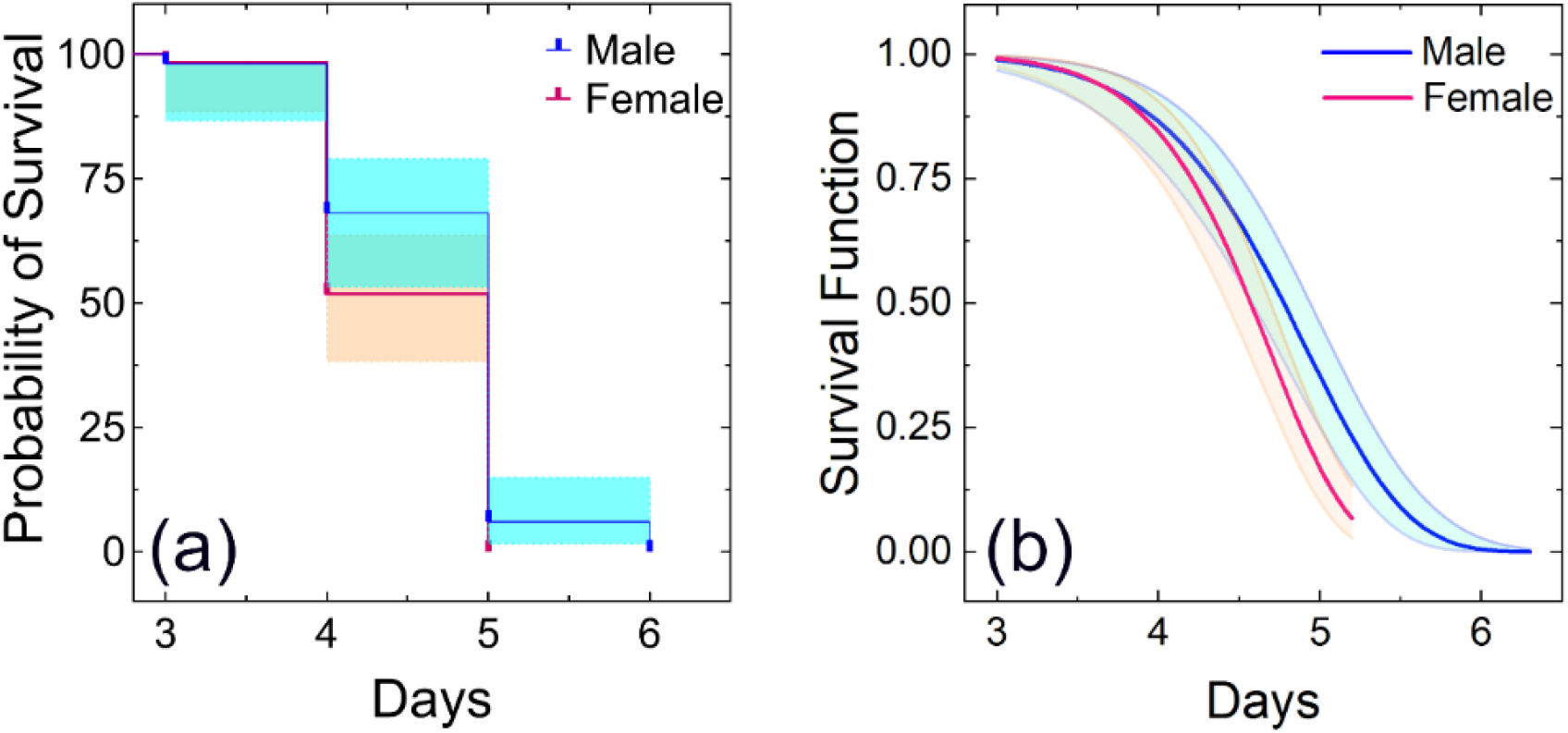
Survival curves of males (n = 50) and females (n = 58) of adult *Bactrocera dorsalis* fed with only distilled water: (a) Survival curves using Kaplan–Meier estimator (b) Survival curves using Weibull parametric model. The color shadows over the survival curves, shown in figure (a) and (b), represent the 95% confidence interval.

A significant difference in mean longevity was observed between adults of two groups: one group was only provided with distilled water, whereas the other group was provided with access to a complete diet. These findings point out the critical influence of diet composition, specifically protein and carbohydrate availability, on adult *B. dorsalis* survival. Furthermore, significant differences in sex-specific longevity patterns were observed under laboratory rearing conditions with access to protein and carbohydrate-rich food, where females exhibited significantly longer lifespans than males, as shown in Figure 6 (a, b). Similar findings were also reported by Jaleel *et al*. and Pieterse *et al*. (Jaleel *et al*., 2017; Pieterse *et al*., 2019).

Welma Pieterse *et al*. reported that the male and female lifespans of *B. dorsalis* were 42.3±25.3 days and 40.03±25.7 days, respectively, when fed a 3:1 mixture of sugar and yeast (Pieterse *et al*., 2019). However, survival analyses in this study showed significantly longer lifespans: 110.26±33.11 days for females and 88.75±28.21 days for males when provided with two types of adult food. These lifespans also exceed those reported by Chen *et al*. across all the tested diets (Chen *et al*., 2013).

The survival analyses, as shown in Figure 7 (a, b), suggest that adults may survive only about 3 to 6 days with only water without any other food sources. In this case, the average longevity of females was approximately 4.50 ± 0.54 days, while the males had a slightly longer average longevity of about 4.72 ± 0.61 days. This survival period is enough for adult flies to disperse over long distances in search of food within the release area.

Overall, this study showed that the survival of *B. dorsalis* varied significantly depending on their food sources and sex. Similar findings were also reported by previous researchers (Chen *et al*., 2013). These results demonstrate that *B. dorsalis* developed well on our formulated larval and adult diets, suggesting significant potential for the use of these formulated diets in mass laboratory-rearing of *B. dorsalis* larvae and adults for Sterile Insect Technique (SIT) program.

## Conclusion

In this study, the effectiveness of our new formulated agar-based larval and yeast-based adult diets for lab-reared *B. dorsalis* was investigated. Statistical analyses including Tukey tests and survival analyses were performed for understanding the efficacy of our formulated diets. Our gel-based larval diet significantly enhanced pupal size (15.18±0.48 mg) and resulted in a high rate of adult emergence (96.88±1.56%). Compared to wild flies, laboratory-reared adults exhibited a notably larger body size, with males measuring 6.77±0.20 mm and females measuring 7.05±0.34 mm. Additionally, our gel-based larval diet formulation reduces waste production by eliminating the bulking agents typically required in larval diets. In addition to larval diet, our protein-based adult diets also exhibited improved parameters related to adult rearing. Adult flies that were fed yeast-based diets produced eggs with a hatching rate of 90.98±3.53% when the eggs were seeded onto the gel-based larval diet. Lab-reared flies using our formulated diet showed superior flight ability (86.392±2.41%), exceeding required mass raring standards. In addition, it was observed that the ovary of lab-reared female flies was larger in length as compared to previous reports, whereas the size of lab-reared male testes remained comparable to wild flies. Furthermore, this study showed that depending on food sources and sex, the survival of *B. dorsalis* varied significantly. While the adults survived only 3 to 6 days on a water diet without food supply, our formulated complete adult diets showed significantly increased longevity (110.26±33.11 days for females, 88.75±28.21 days for males). Importantly, the formulated diets employed for the rearing of larvae and adults in this study adhere to all the minimum quality standards outlined in the FAO/IAEA/USDA quality control manual version 7.0 (FAO/IAEA/USDA, 2019). Moreover, our formulated diets also fulfilled established protocols requirement for mass rearing for the release of sterile males of *B. dorsalis*. The integration of our new formulated diets into the mass rearing process of *B. dorsalis* in a laboratory setting has the potential to significantly improve the production of larger and healthier flies on a large scale, exhibiting improved flight capabilities and reproductive parameters.

## CRediT authorship contribution statement

**Mahfuza Momen:** Conceptualization, Data curation, Formal Analysis, Funding acquisition, Investigation, Methodology, Project administration, Resources, Software, Supervision, Validation, Visualization, Writing original draft, Writing review & editing. **Md. Shahjalal:** Investigation. **Md. Ashikur Rahman:** Investigation. **Md. Aftab Hossain:** Writing review & editing. **Md. Kamruzzaman Munshi:** Investigation. **Kajla Sheheli:** Funding acquisition, Writing review & editing.

## Data availability

The experimental data will be provided upon request.

